# Human Milk Oligosaccharides modulate the risk for preterm birth in a microbiome dependent and independent manner

**DOI:** 10.1101/683714

**Authors:** Manuela-Raluca Pausan, Vassiliki Kolovetsiou-Kreiner, Gesa Lucia Richter, Tobias Madl, Elisabeth Giselbrecht, Eva-Christine Weiss, Evelyn Jantscher-Krenn, Christine Moissl-Eichinger

**Author notes:** These authors contributed equally (Evelyn Jantscher-Krenn, Christine Moissl-Eichinger). E-mail addresses: Manuela-R. Pausan Gesa Lucia Richter Tobias Madl Elisabeth Giselbrecht Eva-Christine Weiss Christine Moissl-Eichinger.

## Abstract

**Background:** Preterm birth is one of the leading causes of neonatal mortality. The causes for spontaneous preterm birth (PTB) are multifactorial and remain often unknown. In this study, we tested the hypothesis that human milk oligosaccharides (HMOs) in blood and urine modulate the maternal urinary and vaginal microbiome and influence the risk for PTB. We analyzed the vaginal and urinary microbiome of a cross-sectional cohort of women with and without preterm labor and correlated our findings with measurements of metabolites and HMOs in urine and blood.

**Results:** We identified several microbial signatures associated with short cervix, PTB and/or preterm contractions such as *Lactobacillus jensenii*, *L. gasseri*, *Ureaplasma sp*. and *Gardnerella sp..* Additionally, we observed associations between sialylated HMOs, in particular 3’-sialyllactose, with PTB, short cervix and increased inflammation and confirmed an influence of HMOs on the microbiome profile.

**Conclusions:** Identifying serum and urinary HMOs and several key microorganisms associated with PTB, our findings point at two distinct processes modulating the risk for PTB. One process seems to be driven by sterile inflammation, characterized by increased concentrations of sialylated HMOs in serum. Another process might be microbiome-mediated, potentially driven by secretor-active HMOs in urine. Our results support current efforts to improve diagnostics and therapeutic strategies.

## Background

Each year, approximately 15 million infants worldwide are born preterm. The rate across 184 countries ranges from 5-18% of all births (Europe: 6%; [1, 2]). Prematurity is the leading cause of mortality in children under the age of five years [1, 3]. Besides the emotional burden on the affected family, preterm birth (PTB) also has severe economic consequences for society. PTB is defined as birth prior to the 37th week of gestation [4], and for unknown reason, the incidence of preterm births is rising.

Spontaneous PTB is a multifactorial process, often accompanied by infections, inflammation, utero-placental ischemia, stress or other immunological mediated processes [5]. Risk factors for PTB include certain lifestyle habits (such as smoking, alcohol consumption, drug abuse, stress), demographic factors (mother age over 35 or under 17, ethnicity, low socioeconomic status), medical conditions (infections, diabetes, high blood pressure, obesity, multiple pregnancies, immunological complications), and genetic predisposition [6]. However, in up to half of all cases, the causes remain unknown [7].

With the current advances to analyze the human microbiome in greater detail, differences in maternal microbiome have been linked to PTB. The vaginal microbiome, which is generally categorized into five community state types (CSTs), undergoes substantial changes over the course of a healthy pregnancy [8]. Diversity and richness of the vaginal microbiome decreases, leading to an enrichment of specific *Lactobacillus* species and a decrease in the relative abundance of *Ureaplasma* and *Mycoplasma* [9]. By lactic acid production, particularly lactobacilli contribute to lowering the vaginal pH, which protects from ascending infections. Perturbation of this well-attuned microbiome, followed by microbial imbalance (dysbiosis) or changes in specific taxa, might lead to infections and PTB [10]. However, the positive or negative role of lactobacilli seems to be rooted on species, or even strain level. On the one hand, high abundance of certain *Lactobacillus* species, but on the other hand, also a generally low abundance of lactobacilli (along with bacterial vaginosis) has been associated with PTB [11][12]. A very recent study, however, highlighted the role of an immune-system component, namely beta-defensin-2, which was found to be negatively correlated with spontaneous PTB, largely independent of the microbiome profile assessed [13]. A causal link between vaginal microbiome composition and PTB has not been established to date.

A potential other risk factor for PTB are urinary tract infections (UTIs). UTIs often remain asymptomatic and hence are overlooked. High-throughput sequencing-based microbiome studies linking urinary microbiome with PTB are rather rare, but a recent study showed that specific microbial taxa, such as *Serratia marcescens* in urine were increased in women with PTB [14].

Recently, human milk oligosaccharides (HMOs) have emerged as potential novel factors involved in maintaining a healthy pregnancy. HMOs, structurally divers and highly bioactive glycans, were found in serum [15] and urine of pregnant women [16, 17] as early as 10 weeks of gestation, and thus, are produced long before birth and start of lactation. In maternal serum, HMO concentrations increase, and composition changes over the course of gestation [15]. As the structure of an HMO determines its function, the relative abundance of individual structures may shape the resulting effect of a complex HMO mixture [18]. High individual variance in HMOs is partly explained by genetic polymorphisms, most importantly of the *Secretor* gene, encoding for the α1-2 fucosyltransferase-2 (FUT2). Individuals with negative secretor status lack FUT2 activity and thus, α1-2 fucosylated structures on cell surfaces in mucosal tissues and bodily fluids. Therefore, the presence of α1-2 fucosylated HMOs such as 2’-fucosyllactose (2’FL) or lacto-N-difucotetraose (LDFT) in pregnant or lactating women is indicative of a positive secretor status. Secretor status, but also metabolic factors such as body composition and BMI were found to be associated with HMO concentration and composition in maternal serum [15], with so far unknown consequences for pregnancy outcomes.

HMOs may mediate their various functions either directly by acting on body cells, or indirectly, by acting on microbial commensals or pathogens. For example, HMOs have been shown to exert immunomodulatory, anti-inflammatory effects for example by regulating the expression of macrophage derived cytokines, thereby inducing Th2 polarization or dampening LPS response [19, 20]. On the other hand, HMOs can shape microbial communities as shown for the infant’s gut. HMOs were found to serve as substrates for specific strains of genera such as *Bifidobacterium* and *Bacteroides* [21, 22], and *Lactobacillus* [23, 24]. Besides their prebiotic activities towards beneficial microbes, HMOs can also directly act on microbes as anti-adhesives [25–28] or antimicrobials [29]. These findings indicate that HMOs present at particular body sites might have direct or indirect effects on microbial communities at these sites. Although the relatively high abundance of specific HMOs in urine has been known for decades [16], to date, the physiological composition or biological roles of urinary HMOs in pregnancy related disorders such as PTB have not been investigated.

Here, we hypothesize that the maternal microbiome in urine and vagina plays a critical role in triggering PTB. We also consider HMOs in serum and urine as important regulators of the microbiome, which might cause subtle changes in certain microbial taxa, preventing or promoting PTB. To test this hypothesis, we performed a cross-sectional study to understand the interplay of the microbiome profile (catheter urine and vagina), the urinary metabolome and the serum and urine HMOs in women with preterm labor and thus at risk for spontaneous PTB (n=33) and compared the results with those of appropriate controls (n=27).

## Results

A total of 60 women were recruited during examination at the hospital due to potential preterm contractions. All women included were between weeks 23-34 of pregnancy. Before any treatment, a vaginal swab, catheter urine and blood were collected from all women.

After medical examination and transvaginal scan for cervical length measurements, around 55% of the women received tocolysis to prevent PTB. Tocolysis is recommended in the case of regular preterm contractions (more than four contractions within 20 minutes) and cervical shortening of below 25 mm, assessed by transvaginal ultrasound.

Tocolysis and control group were analyzed with respect to their vaginal and urinary microbiome (qualitative), the bacterial load (quantitative), urine metabolites, and HMOs (blood and urine). All metadata (Suppl. Table 1) were used for correlation analysis. Socio-demographic characteristics of the two groups of women are summarized in Table 1. 40% of the women had a short cervix (< 25 mm) at the moment of the recruitment. The majority of these women received tocolysis (“cases”). Cases did not differ significantly from the control group in ethnicity, body mass index, and age (Table 1). 18.3% of all recruited women delivered preterm, while the majority of the women delivered after 37 weeks of pregnancy. Except for two, all women with PTB had a short cervix at presentation at the clinics (81.8%). The variance in cervical length was also found to be the only statistically significant difference between case and control groups. Not all women who delivered preterm received tocolytic treatment (Table 1).

**Table 1:** Group characteristics

|  | TOCOLYSIS GROUP | CONTROLS | P-VALUE |
| --- | --- | --- | --- |
| RECEIVED TOCOLYTICS | yes (n=33) | no (n=27) |  |
| AGE | 29.45 ± 0.895 (20-39) | 29 ± 1.089 (18-41) | 0.746 |
| 18-25 | 7 (21.21%) | 6 (22.22%) |  |
| 26-35 | 21 (63.64%) | 16 (59.26%) |  |
| 36-41 | 5 (15.15%) | 5 (18.52%) |  |
| CERVICAL-LENGTH | 22.45 ± 1.6504 (6-40) | 34.92 ± 1.90 (6.8-50) | 0.000* |
| BMI | 22.27 ± 3.56 (15.43-30.86) | 24.31 ± 4.5 (16.94-35.66) | 0.055 |
| UNDERWEIGHT (<18,5) | 5 (15.15%) | 4 (14.82%) |  |
| NORMAL WEIGHT (18,51-24,9) | 22 (66.67%) | 11 (40.74%) |  |
| OVERWEIGHT (25,0-29,9) | 4 (12.12%) | 9 (33.33%) |  |
| OBESE (>30) | 2 (6.06%) | 3 (11.11%) |  |
| ETHNICITY |  |  | 0.141 |
| WHITE | 32 (97%) | 23 (85.2%) |  |
| ASIAN | 0 (0.0%) | 3 (11.1%) |  |
| BLACK | 1 (3%) | 1 (3.7%) |  |
| GESTATION AT SAMPLE (WEEKS) | 29<sup>+2</sup> ± 3,3 (23<sup>+3</sup> – 33<sup>+4</sup>) | 28<sup>+4</sup> ± 3,2 (23<sup>+4</sup> – 33<sup>+1</sup>) | 0.659 |
| DELIVERY |  |  |  |
| TERM | n=24 (72.7%) | n=25 (92.6%) | 0.091 |
| PRETERM | n=9 (27.3%) | n=2 (7.4%) |  |
| CRP | 7.43 ± 1.3217 (0.9-30.1) | 9.763 ± 3.2984 (0.8-86.2) | 0.841 |
| SECRETOR STATUS |  |  |  |
| SECRETOR | n=29 (87.9%) | n=21 (77.8%) | 0.322 |
| NON-SECRETOR | n=4 (12.1%) | n=6 (22.2%) |  |
| VAGINAL PH | 4.85 ± 0.1418 (4-7) | 4.73 ± 0.1623 (3-7) | 0.743 |
| URINE PH | 6.53 ± 0.1442 (5.5–8) | 6.69 ± 0.311 (5-8) | 0.213 |

\*p-value < 0.05

### HMO signatures in serum and urine are different in women prone to PTB

We detected HMOs in all samples of maternal serum and urine and quantified the five most abundant HMOs using a HPLC-based method. In serum, median concentrations (95% confidence interval [CI] of median) for the five most abundant serum HMOs, were 0.68 nmol/mL (95% CI of median 0.41-0.81) for 2’FL, 0.23 nmol/mL for LDFT (95% CI of median 0.16–0.36), 0.37 nmol/mL (95% CI of median 0.32-0.41) for 3’SL, 0.093 nmol/mL (95% CI of median 0.087-0.10) for 3’SLN and 0.24 nmol/mL (95% CI of median 0.21-0.27) for 6’SLN (Fig. 1a). The same HMOs species were present in urinary samples, although total concentration was approximately 50-fold higher (Fig. 1a). Urine profiles also showed a significantly higher proportion of the fucosylated HMOs 2’FL, LDFT and lower proportion of the sialylated HMOs, 3’SL, and 6’SLN (Fig. 1b). The secretor active HMOs, 2’FL and LDFT, were strongly correlated between serum and urine samples. In 10 out of 60 women, we detected only trace amount of 2’FL and LDFT in serum and urine samples, indicating secretor negative status (Fig. 1c). The sialylated HMOs 3’SL and 3’SLN in serum and urine were also correlated, but to a lesser extent (Fig. 1c). Normalization of urinary HMOs with the osmolality resulted in a marked increase in R squared values of the correlations between serum and urine for 2’FL, LDFT, and 3’SL (Fig. 1c).

**Fig. 1.**
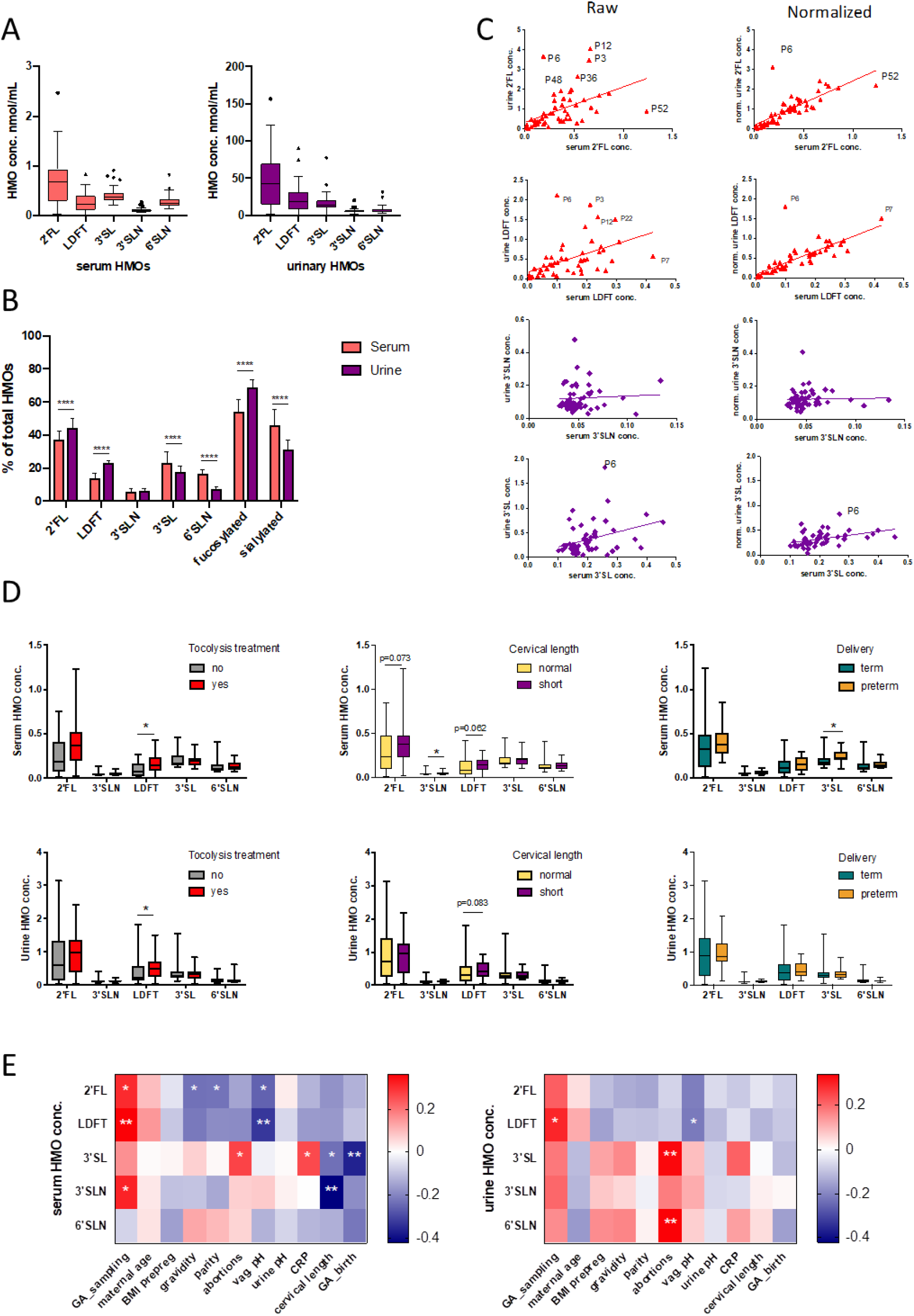
Serum and urinary HMOs are associated with preterm indicators and PTB. **a** Absolute HMOs concentration in serum (red boxes) and in urine (purple boxes). **b** Relative HMO concentrations of serum and urine. **c** Linear regression of serum and urine 2’FL (first row), LDFT (second row), 3’SLN (third row) and 3’SL (forth row) before and after normalization with osmolality. **d** Specific HMO concentrations were different in serum (upper row) and urine (lower row) within the three groups (Mann-Whitney t-test). LDFT was increased in pregnant women who received tocolytic treatment compared to controls; 3’SLN was higher in women with short cervix versus normal cervix and 3’SL was increased in women who delivered preterm versus women with term deliveries. **e** Heat map of Spearman correlations between HMO concentration and maternal variables. Serum 3’SL was positively correlated with CRP, and negatively with cervical length and gestational age (GA) at birth. 2’FL, 2’-fucosyllactose; LDFT, Lacto-N-difucotetraose; 3’SL, 3’-sialyllactose; 3’SLN, 3’-Sialyllacosamine; 6’SLN, 6’-sialyllactosamine; *p <0.05; **p < 0.01; ***p < 0.001; **** p < 0.0001.

To test whether HMO concentrations were different in women with and without labor, we compared HMOs in serum and urine between cases and controls and found significantly higher LDFT concentrations in the cases (Mann Whitney t-test) (Fig. 1d). Two women in the control group who did not receive tocolytic treatment, delivered their babies preterm three and four days after admission to the clinics. Excluding them from the analysis did not change results. In women with a short cervix, we found significantly higher 3’SLN and a trend towards higher 2’FL and LDFT in serum compared to women with normal cervix. In urine, there was a trend towards higher LDFT in women with short cervix without reaching significance (Fig. 1d). Notably, serum 3’SL was significantly higher in women with subsequent PTB compared to women who delivered at term, whereas urinary HMOs were not altered with PTB (Fig. 1d).

We next performed Spearman correlation analyses to assess whether HMOs were associated with cervical length, gestational age at birth, CRP (as marker of inflammation) and maternal anthropometrics data (Fig. 1e). All HMOs, especially serum LDFT and 2’FL were positively correlated with gestational age at the admission at the clinics (Spearman rho r=0.36, p=0.004; r=0.3, p=0.015, respectively). Serum 3’SL was negatively correlated with gestational days at birth (r=-0.37, p=0.004), and with cervical length (r=-0.23, p=0.045), and positively correlated with CRP levels (r=0.26; p=0.032). Serum 3’SLN was negatively correlated with cervical length (r=-0.42, p=0.001). Serum 2’FL and LDFT were significantly negatively correlated with vaginal pH (r=-0.26, p=0.032; r=-0.22, p=0.009, respectively), whereas associations with urinary 2’FL and LDFT did not reach significance (Fig. 1e). When we controlled for the timing of sampling, all the above associations remained significant.

### Microbial density and richness in vaginal and urinary samples is not indicative of preterm labor

We next used the catheter urine and vaginal swabs to assess differences in the microbial diversity and density related to preterm labor, short cervix and PTB. The qPCR results showed no significant differences in the number of 16S rRNA gene copies in neither urine nor vagina, within the tocolytic group (no/yes), cervical length group (normal/short) or delivery group (term/preterm) (Suppl. Fig. S1).

When comparing the vaginal with the urinary microbial composition, we found a generally higher microbial load in the vaginal samples (Fig. 2c). The number of bacterial 16S rRNA gene copies in urine varied between 10^2^ and 10^7^ 16S rRNA gene copies/mL, being in agreement with previously published data [30]. In general, the urinary microbiome was more diverse and richer based on the results from alpha diversity measurements (Chao1, Shannon, and inverse Simpson) shown in Fig. 2a.

**Fig. 2.**
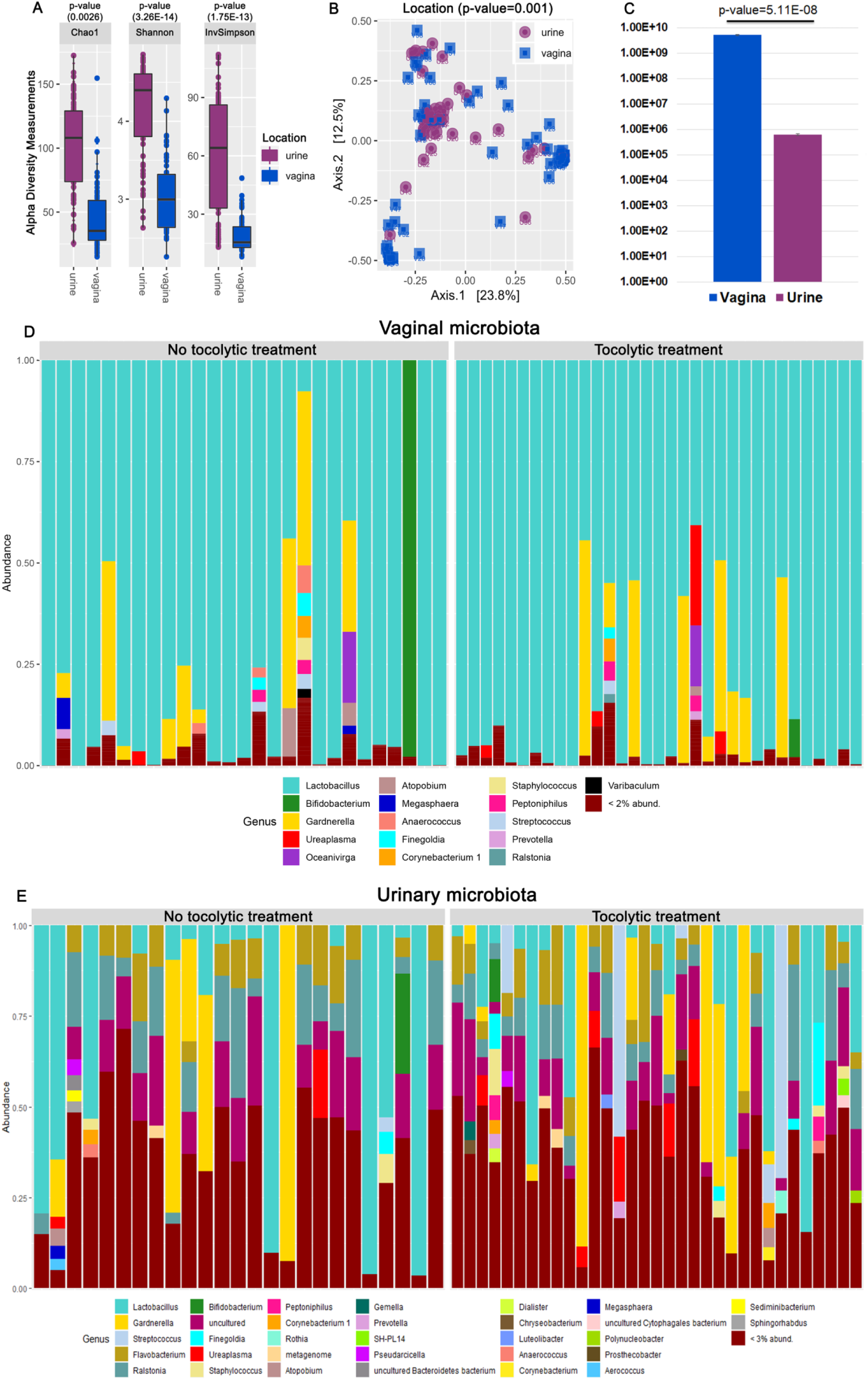
The urinary and vaginal microbiome: alpha, beta diversity and microbial composition. **a** Alpha diversity of the two body sites shows that the urinary microbiome is richer and more diverse compared to the vaginal microbiome. **b** Beta diversity based on Bray-Curtis dissimilarities show that the urine microbial community is different than the vaginal microbial community, although the two do not cluster entirely separately. **c** The number of 16S rRNA gene copies (qPCR results) in urine is lower than in the vagina. **d** The barplot depicts the microbial composition of vagina at genus level showing a dominance of *Lactobacillus* species. **e** The barplot shows the urine microbiota at genus level being dominated by several different genera such as: *Lactobacillus*, *Gardnerella, Streptococcus* and *Flavobacterium*.

Furthermore, the beta diversity (Fig. 2b) indicated that the urinary and vaginal microbial communities were distinct (p=0.001), although both shared some similarities like the presence of *Lactobacillus*, *Prevotella*, *Gardnerella, Ureaplasma, Anaerococcus* and other microbial species (see below).

### Certain Lactobacillus, Gardnerella and Ureaplasma signatures in the vaginal microbiome, but not necessarily community state types, correlate with preterm indicators and PTB

In our study, the vaginal microbiome was dominated by *Lactobacillus*, followed by *Gardnerella*, *Atopobium*, *Ureaplasma*, *Anerococcus*, *Finegoldia* and other genera (Fig. 2d), and thus, was similar to results from previous studies [10, 31].

Performing hierarchical clustering at species level, we classified the vaginal samples into five CSTs [10]) (Fig. 3a), which are characterized as follows: CST I (*L. crispatus*), CST II (*L. gasseri*), CST III (*L. iners*), CST IV (diverse species), and CST V (*L. jensenii*). The most prevalent CST observed in our cohort was CST I (30/60, 50%), followed by CST III (14/60, 23.3%), CST IV (9/60, 15%), CST II (4/60, 6.7%) and CST V (3/60, 5%).

**Fig. 3.**
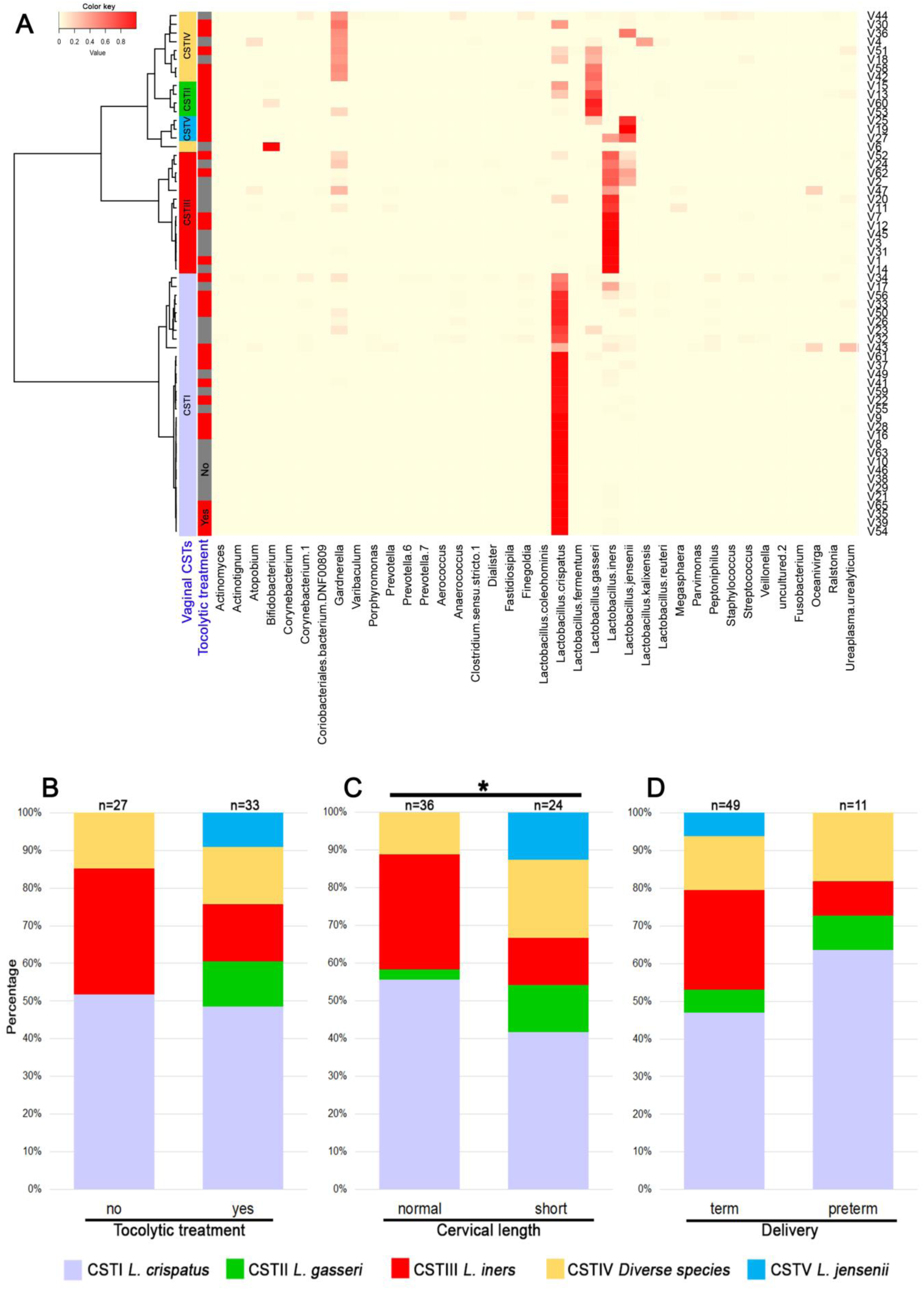
Vaginal microbiome clustered in community state types (CSTs), and prevalence of CSTs in different groups. **a** Community state types as found in our cohort (heat map). **b**, **c, d** The barplots show the community state types (CSTs) and their prevalence in the **b** tocolytic treatment group, **c** cervical length group and **d** delivery group.

CST II and CST V were only present in cases, and mainly in those with a short cervix (< 25 mm), but not in the subjects who delivered preterm (Fig. 3b, c, d), indicating an association between *L. jensenii*/ *L. gasseri* predominance and preterm contractions and short cervix. We found significant differences in the incidence of different CSTs in women with normal versus short cervix, but no differences in the groups formed by either tocolytic treatment (no, yes) or delivery (term, preterm) (Fig. 3b, c, d).

Analyzing alpha (Chao 1, Shannon and inverse Simpson) and beta (principal coordinate analysis, PCoA) diversity for the vaginal microbiome in the three groups (tocolytic treatment, cervical length and delivery outcome), we found no significant differences with respect to diversity, richness and clustering, indicating similar microbial communities within the groups (Suppl. Fig. S2a-f).

To further investigate associations between specific taxa and preterm contractions, cervical length or PTB, we performed LEfSe (Linear discriminant analysis effect size) for all three groups. Certain ribosomal sequence variants (RSVs) affiliated with *Lactobacillus jensenii* were indicative for preterm contractions (RSV31: p=0.038, ANOVA) and short cervix (p=0.019), but not with PTB (p=0.31), while *Lactobacillus iners* was associated with normal cervix (RSV13: p=0.041, ANOVA) and RSVs of *Lactobacillus crispatus* with PTB. *Ureaplasma urealyticum* RSVs were correlated with PTB and preterm contractions (Fig. 4a, b, c).

Additionally, we investigated associations between taxa (Fig. 4d) or genera (Suppl. Fig. S2g) with retrieved cohort characteristics performing Spearman correlation analysis and observed similar correlation patterns. RSVs of *Lactobacillus jensenii* and *Lactobacillus gasseri* strongly correlated with preterm contractions and short cervix. We found similar correlations between certain *Gardnerella* species and preterm contractions, short cervix, and PTB, while other *Gardnerella* species were correlated with term delivery. At genus level, certain genera such as *Staphylococcus*, *Ureaplasma*, *Streptococcus* and *Finegoldia* correlated with PTB. *Ureaplasma* was also correlated with tocolytic treatment, but not with short cervix (Suppl. Fig. S2g).

**Fig. 4.**
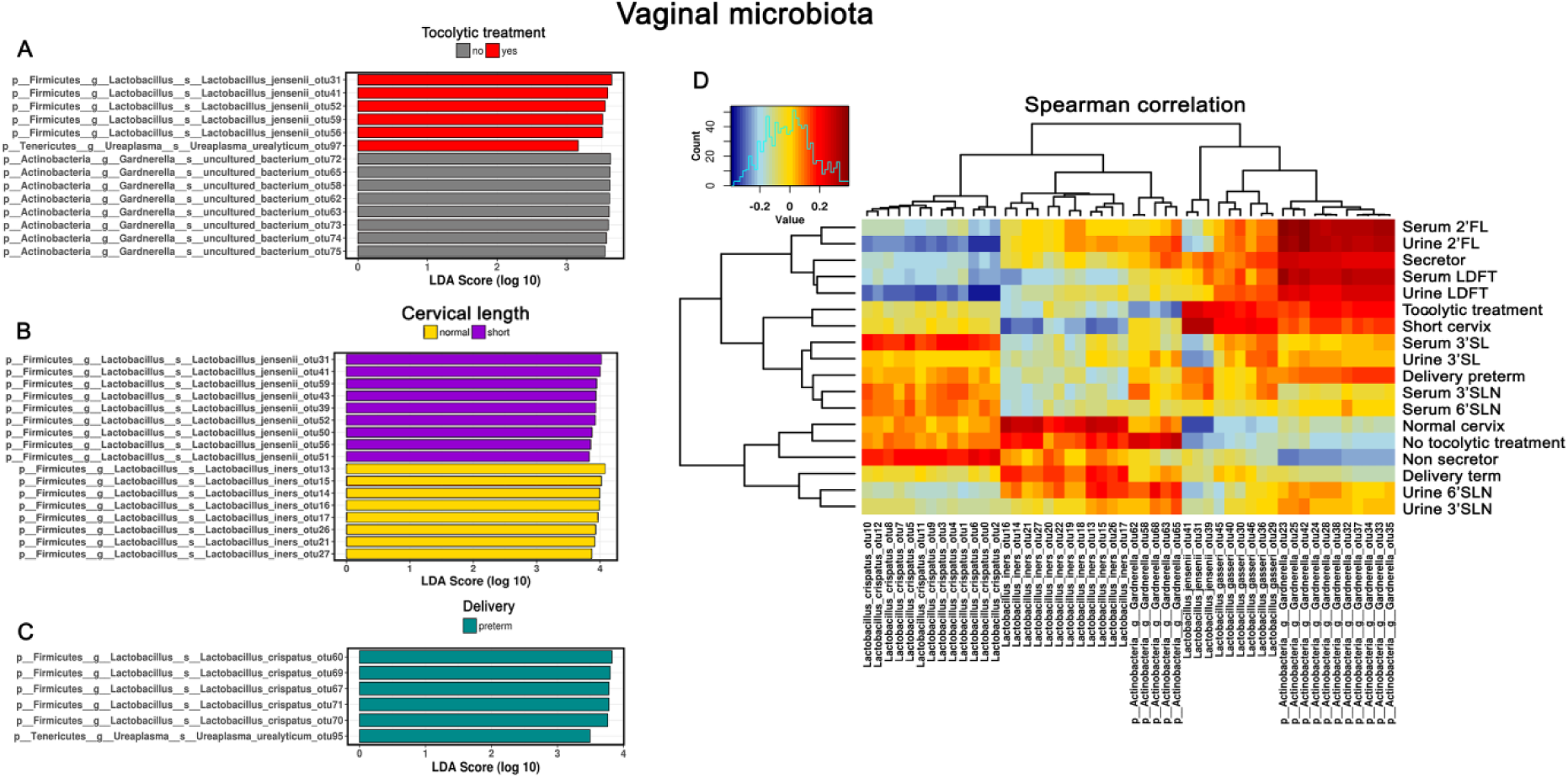
LEfSe analysis and Spearman correlation heat maps of vaginal microbiome. **a, b, c** LEfSe (Linear discriminant analysis effect size) of vaginal microbiome indicate the association of specific lactobacilli with **a** preterm labor, **b** cervical length and **c** delivery outcome. **d** Spearman correlation heat map of the same data (RSV level), shows different associations between vaginal microbial signatures and different characteristics, such as HMOs, delivery outcome, tocolytic treatment etc.

### Microbiome signatures associated with vulnerability to PTB are also detectable in urinary samples

The microbiome in cathetered urine was very diverse and rich compared to vaginal microbiome. The microbial community included signatures of species belonging to *Lactobacillus*, *Gardnerella*, *Streptococcus*, *Dialister*, *Finegoldia*, *Ureaplasma*, *Staphylococcus*, *Atopobium*, Anaerococcus, *Peptoniphilus* and many more (Fig. 2e), also identified in previous studies as part of the normal urinary microbial community [32].

To explore differences in the urinary microbial communities within the groups formed by the tocolytic treatment, cervical length and time of birth, we analyzed alpha and beta diversity using Chao1, Shannon, and inverse Simpson indices. The results showed no differences in the alpha diversity within the three groups analyzed (Suppl. Fig. S3a-f). PCoA based on Bray-Curtis dissimilarities and Adonis tests also revealed no significant differences, indicating similar microbial communities in urine (Suppl. Fig. S3d, e, f).

LEfSe analysis identified several taxa to discriminate between the analyzed groups. RSVs of *Lactobacillus jensenii* (RSV41: p=0.021, ANOVA), *Lactobacillus gasseri* and species of *Finegoldia* (RSV102: p=0.043, ANOVA) were associated with short cervix, while species of *Pseudarcicella* were associated with normal cervix. *Lactobacillus crispatus* was correlated with term deliveries and women who did not receive tocolytics (RSV0: p=0.041, ANOVA), while *Ureaplasma urealyticum* was associated with PTB (RSV100: p=0.051, ANOVA) (Fig. 5a, b, c). Spearman correlation analyses between the urinary microbiome and variables of the three groups revealed similar associations. In addition, specific species of *Gardnerella* were correlated with PTB, short cervix, preterm contractions and all five HMOs investigated in serum and urine (Fig. 5d). At genus level, *Gardnerella* was positively correlated with all measured HMOs, PTB, short cervix, and tocolytic treatment. *Ureaplasma* was mainly associated with PTB and Non-secretor status, and *Pseudarcicella* was positively correlated with normal cervix (Suppl. Fig. S3g).

**Fig. 5.**
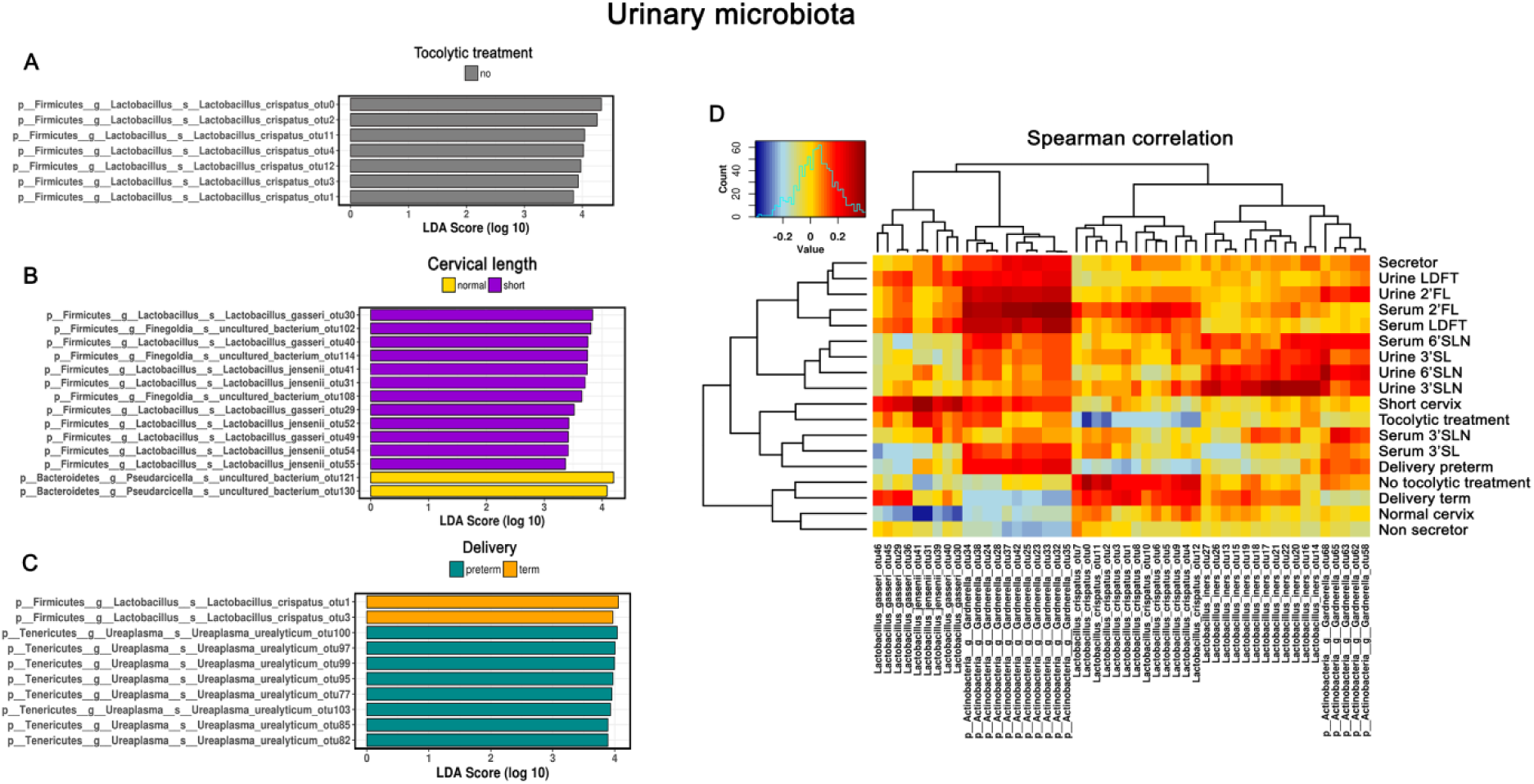
LEfSe analysis and Spearman correlation heat maps of urinary microbiome. **a, b, c** LEfSe (Linear discriminant analysis effect size) of urinary microbiome indicate associations of specific lactobacilli with **a** preterm labor, **b** cervical length and **c** delivery outcome. **d** The Spearman correlation heat map shows similar associations between specific *Lactobacillus* species and preterm labor, cervical length and delivery outcome, furthermore *Gardnerella* species were also associated with specific HMOs and preterm delivery

### The urinary metabolites are unaffected by preterm labor

Principal Component Analysis (PCA) plots of urinary metabolites revealed no differences between women with short cervix and normal cervix (Suppl. Fig. S4), or women with term delivery and preterm delivery (Suppl. Fig. S5). Women treated with tocolytics had slightly decreased urinary citrate concentration compared to controls (Suppl. Fig. S6c), but no differences in other metabolites (Suppl. Fig. S6). We also compared the metabolites acetate, trimethylamine and tyrosine, which have been previously associated with PTB [33, 34], but found no significant differences between cases and controls (Suppl. Fig. S7).

### The microbiome in urine and vagina follows the HMO profile in serum and urine

Having found differences in HMOs and certain microbial taxa related to preterm indicators and PTB, we asked whether HMOs and microbiome are linked. Indeed, we found strong correlations of HMOs in serum and urine with urinary and vaginal microbiome composition. Serum and urinary HMO concentrations were strongly correlated with *Gardnerella* species in the urinary and vaginal microbiome. Specific vaginal genera such as *Ureaplasma*, *Gardnerella*, and *Flavobacterium* were positively correlated with the concentrations of LDFT and 2’FL in both, urine and blood, and thus, with positive secretor status (Suppl. Fig. S2g). Vaginal *Flavobacterium* genus was correlated with sialylated HMO concentrations in urine and with serum 3’SLN (Suppl. Fig. S2g). Vaginal *Ureaplasma* was correlated with serum 6’SLN (Suppl. Fig. S3G), while the *Ureaplasma* in urine was correlated with sialylated HMO concentrations with the exception of serum 3’SLN. Serum 3’SLN was mainly correlated with *Finegoldia* (Suppl. Fig. S3g).

We also observed correlations between *Bifidobacterium* in urine and 6’SLN and 3’SL in urine and serum (Suppl. Fig. S3g). Furthermore, we found correlations between specific HMOs and *Lactobacillus iners* from urine samples (Fig. 5d). *Gardnerella* species were also highly correlated with LDFT and 2’FL concentrations in serum and urine, and thus, with positive secretor status. *Lactobacillus crispatus* species were highly positively correlated with serum 3’SL concentration.

### Inflammation, either directly mediated by sialylated HMOs or indirectly through affecting the microbiome, might increase the risk for PTB

In a next step, we were interested in the microbial functions and their correlation with HMOs and other obtained metadata. For this, we predicted functional profiles from our 16S rRNA gene data using Tax4Fun. As shown in Suppl. Fig. S8, the functional profiles of the urinary microbiome grouped into three major groups. Notably, one cluster of microbial functions was strongly influenced by the secretor status, reflected by an increase in sugar-degradation-related microbial functions (e.g. pentose and glucoronate interconversions (p=0.031, ANOVA), pentosephosphate pathway, galactose metabolism, starch and sucrose metabolism). This observation explains the above mentioned increase of particularly sugar-metabolizing microbial signatures, such as *Gardnerella,* along with secretor-associated HMOs LDFT and 2’FL.

Notably, microbiome functional profiles in vagina and urine were not correlated with inflammation status (CRP), but microbial function profiles of urine, but not the vagina, were strongly correlated with the number of 16S rRNA gene copy numbers in the urine samples therein (two categories of copy numbers groups, high and low, cutoff-10^5^; p=0.001). These copy numbers were increased along with again sugar-degrading microbial functions (galactose metabolism, starch and sucrose metabolism; LEfSe analysis), and corresponding microbial taxa, such as *Gardnerella* (p=0.029), *Lactobacillus* (p=0.00064) and *Finegoldia* (p=0.0058, ANOVA; Suppl. Fig. S9), indicating an increased microbial load in the urine due to the secretor-associated HMOs. This was reflected also in the vaginal microbiome, as a high copy number in the urine correlated with an increase in *Gardnerella* and *Finegoldia* therein as well.

In Fig. 6, we present an overview on the results of the study, in which we were able to identify potential key compounds and key microorganisms and their interplay modulating the risk for PTB (Fig. 6a). Sialylated HMOs, in particular 3’SL, were positively correlated with PTB, short cervix, and increased inflammation. Notably, 3’SL only partially correlated with key microbial signatures in urine and vagina, indicating an independent risk increase through a direct, but microbiome-independent mechanism (Fig. 5B). Key microbial signatures were *Lactobacillus jensenii*, *L. gasseri*, *Ureaplasma* sp., *Gardnerella* sp. and *Finegoldia* sp., which appeared to be correlating with short cervix, PTB and/or preterm contractions. Notably, most of the above mentioned microorganisms were associated with an increased bacterial load in the urine, indicating a potential low-inflammatory or beginning infection therein. Particularly, *Gardnerella* signatures were found to be associated with the secretor status, in agreement with the observations on functional level. The secretor status, however, was also found to be associated with a higher bacterial load in the urine.

**Fig. 6.**
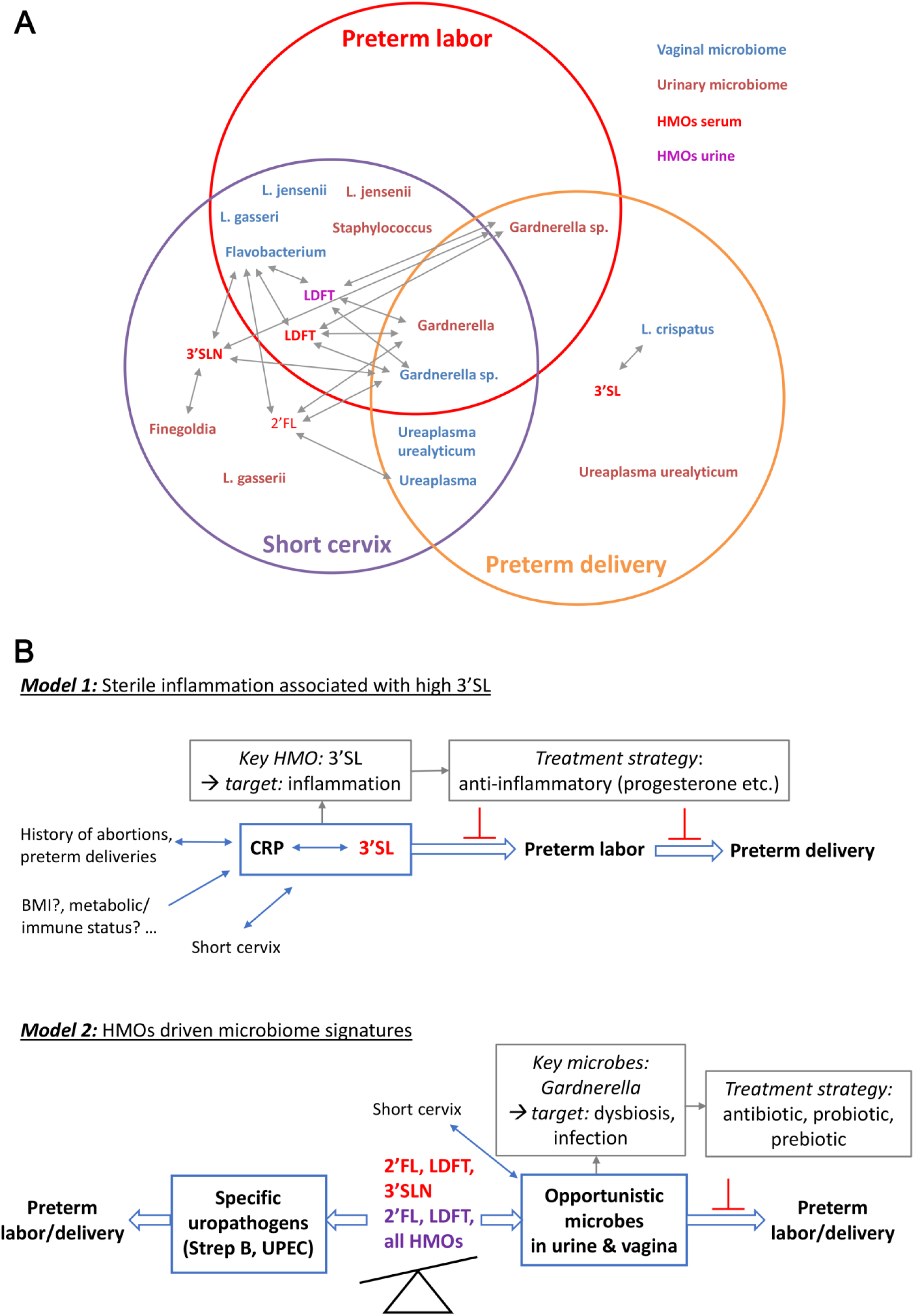
Overview on the results and the proposed models. **a** Venn diagram shows specific HMOs (red for serum, purple for urinary) and microbial genera or species (blue for vaginal and brown color for urinary) that were found associated with preterm labor, short cervix or preterm delivery, and their interplay. Arrows indicate correlations between specific HMOs and microbes. **b** Two distinct models on how HMOs might be involved in modulating the risk for preterm birth and potential treatment strategies are shown. Model 1 is characterized by an increased concentration of sialylated HMOs, i.e. 3’SL, associated with sterile inflammation, independent of the vaginal or urinary microbiome. Thus, anti-inflammatory therapies might be recommended. Model 2 is characterized by an imbalance in HMOs, leading to disturbances in vaginal and urinary microbiome. This, combined with other risk factors (short cervix), might promote preterm labor and delivery. While specific HMOs might prevent colonization with urogenital pathogens, the same HMOs might also foster overgrowth of certain opportunistic microbial species in the urinary tract and then vagina leading to beginning infections and promoting preterm labor and delivery. 2’FL, 2’-fucosyllactose; LDFT, Lacto-N-difucotetraose; 3’SL, 3’-sialyllactose; 3’SLN, 3’-Sialyllacosamine.

To summarize, our observations point at two different processes. One process appears to be driven by sterile inflammation, characterized by increased sialylated HMO concentrations in serum, another process appears to be indirect and microbiome-mediated, which could, however, be driven by secretor-associated HMOs. HMOs thus were identified as key molecules involved in the regulation of term birth (Fig. 6b).

## Discussion

Our study combines the comparative analysis of vaginal and urinary microbiome, serum and urinary HMOs, and urinary metabolites in pregnancies at risk for spontaneous PTB compared with healthy controls. We identified key HMOs and microbial signatures associated with a higher risk of PTB and postulate a key role of HMOs in the regulation of a healthy, term pregnancy.

Few studies have investigated changes in HMO profiles in human milk as a consequence of PTB [35, 36], while their potential causal role in PTB has not been explored so far. Recently, we have shown that HMOs in serum increase during pregnancy, raising the question whether HMOs are associated with preterm labor and/or PTB. An early study reporting on HMOs in urine of pregnant women speculated about the importance of HMOs in maintaining a healthy pregnancy [16]. Herein, we were able to confirm this hypothesis, as specific HMOs were identified, indicating a risk for PTB. Furthermore, we report for the first time, associations of serum and urinary HMOs with specific genera and species of the urinary and vaginal microbiome.

In this cross-sectional study, the sampling window was from 23^+0^ to 34^+0^ weeks of gestation and especially secretor active HMOs strongly varied. In all women, assigned secretor status (based on the relative abundance of 2’FL) matched between serum and urine samples. A secretor negative status was identified in 16% of all women, which was comparable to our previous findings [15]. Correcting for time of sampling and excluding the secretor negative women, we found high serum 3’SLN associated in women with short cervix and high 3’SL associated in women who went on to later deliver preterm. Interestingly, only 3’SLN and 3’SL in serum, and not in urine, were associated with short cervix or PTB. While the discrepancies between serum and urine could be multiple, e.g. attributed to the relatively small sample size, or errors in normalization of urine, this might also indicate a more systemic mechanism.

Our finding that sialylated HMOs in serum are associated with short cervix and PTB are in accordance with a study reporting increased total serum sialic acid concentration associated with PTB [37]. In human milk, HMOs were also found to be different in sialylation after PTB. These findings might support the hypothesis that sialyation, in general, is altered in preterm labor mediated by (sterile) inflammatory processes. This pathway would not necessarily be dependent on changes in urinary and vaginal microbiome and would explain why urinary HMOs are not associated with preterm birth in our cohort excluding clinically manifest infections. In addition, accumulating evidence points towards increased plasma sialic acid in inflammatory pathologies such as cancer, cardiovascular diseases and type 2 diabetes [38]. Similarly, inflammation was shown to be associated with altered glycosylation due to extrinsic sialylation via extracellular sialyltransferases [39]. Whether increased sialylation is a general response to inflammatory conditions in pregnancy, and whether these changes in sialylation have ameliorating or deteriorating effects on the pregnancy remains to be elucidated. Whether these sialylated HMOs in serum might have a potential as predictive markers for PTB also needs to be investigated in accordingly designed larger prospective studies.

Our study aimed to investigate medically unexplained preterm labor. Thus, we excluded women who were diagnosed with or had previously been treated for UTI, which is causally linked to preterm labor and PTB. Previous in vitro studies have shown that HMOs protect against uropathogenic *Escherichia coli* invasion and cytotoxicity [40] or induce growth defects in GBS [41], both common uropathogens associated with UTI. A study showed that negative secretor status was associated with higher incidence of UTI in pregnant women [42], indicating that secretor active structures in the urothelial site might protect from UTI. This seems partially contradictory to our finding that positive secretor status was associated with increased bacterial load in the urine, correlating with an increase of *Gardnerella* and *Ureaplasma* signatures. However, as answering this question was not scope of this study, we excluded symptomatic UTIs. This might have biased the collective, leading to a smaller proportion of secretor negative women at risk for PTB than normally found in the general population. Our results indicate that HMOs in urine (and serum) shape the microbial community, supporting in particular those that are able to feed on them, such as *Gardnerella*. Like bifidobacteria, *Gardnerella vaginalis* possess the fructose-6-phosphate phosphoketolase (F6PPK), the key enzyme of the so-called “bifid shunt”, a specific pathway for carbohydrate metabolism [43, 44]. In accordance with our estimation of the function capacities, this indicates that *Gardnerella* is capable of utilizing HMOs, in particular those that are associated with positive secretor status. This leads us to speculate about a potential adaptation and growth advantage of *Gardnerella* in secretor positive, pregnant women, opening up new questions on longitudinal associations with gestation and increasing HMO concentrations.

Previous studies have reported a decrease in vaginal microbial diversity and richness between first and second trimester in women with PTB [45], while other studies could not identify a change in the microbial diversity and richness [11, 31, 46]. Moreover, in our study, we did not observe significant changes in diversity, richness or bacterial load in women at increased risk. We identified CST V, dominated by *Lactobacillus jensenii,* mainly in women who were at high risk of PTB and received tocolytic treatment, and especially in women who had a short cervix. Interestingly, none of the women who delivered preterm had CST V at the time of sampling. Additionally, we observed an increase in the relative abundance in specific taxa, such as *L. jensenii* and *Ureaplasma urealyticum,* in women who received tocolytic treatment and in women with short cervix. In our study, we also found species of *Gardnerella* associated with preterm labor and short cervix. Others have shown associations between either specific taxa or vaginal CSTs and PTB [8, 12, 46]. The observed associations of CST IV with PTB might be explained by a higher risk for bacterial vaginosis in these women compared to women with a vaginal microbiome dominated by *Lactobacillus* species.

Kindinger et al. [11] showed associations between *Lactobacillus iners*, short cervix and PTB. In our study, *Lactobacillus iners* was not associated with short cervix or PTB. In contrast, more women who delivered term, had a normal cervix and received no tocolytic treatment had a vaginal microbiome dominated by *Lactobacillus iners*. This discrepancy might be explained by differences in the methodology used and the ethnicity of the cohorts. The study cohort of Kindinger et al. included a higher proportion of women of African origin than ours, and 32% of the preterm deliveries were of mothers of African origin. In our study, we had only 2 (3.33%) women of African origin, none of whom delivered preterm. Furthermore, it has previously been shown that vaginal microbiome of African American women is more likely to be colonized by *L. iners* (CST III), *Anaerococcus* or *Mycoplasma* (CST IV), while European women are more likely to be colonized by *L. crispatus*, *L. jensenii* and *L. gasseri* [47]. Therefore, the association found in previous studies between *L. iners* and PTB might be cohort specific.

Our findings that *L. jensenii* and *Gardnerella* abundance in the vaginal microbiome was associated with higher risk of PTB are in accordance with previous studies [46, 48]. However, the observation that vaginal microbiome dominated by *L. crispatus* might be protective against PTB [11, 46, 48] was not confirmed in our study. In contrast, seven out of eleven women who delivered preterm had a vaginal microbiome dominated by *L. crispatus*. We are aware that the number of women who delivered preterm in our cohort is relatively small compared to the other studies and a large cohort of patients delivering preterm is needed to validate previous results.

Secretor status, as assigned by abundance of 2’FL was not associated with preterm labor or PTB, which might due to relatively small sample size. However, CST V (*L. jensenii*) which was associated with preterm labor and short cervix was only found in secretor positive women. None of the secretor negative women (10 out of 60), had CST IV or CST V.

Our results on the urinary microbiome are consistent with a previous study, where no distinct changes were observed in the alpha and beta diversity of urinary microbiome between women who delivered preterm compared to women who delivered term [14]. However, we observed enriched taxa between the groups, with e.g. *L. crispatus* having a higher relative abundance in pregnant women who did not receive tocolytic treatment and especially in patients who delivered term. *L. gasseri, L. jensenii* and species of *Finegoldia* were enriched in women with a short cervix, while *Ureaplasma urealyticum* was increased in women delivering preterm. We also observed strong correlations between PTB, short cervix and species of *Gardnerella*. As none of the women showed symptoms of urinary infections at the time of sampling, the increase in *Ureaplasma urealyticum* in the PTB group speaks for asymptomatic infections which could be monitored as a risk factor. Different to our study, Ollberding et al. have shown an increase in the relative abundance of *Prevotella*, *Sutterella*, *L. iners*, *Blautia*, *Kocuria*, *Lachnospiraceae* and *S. marcescens*, which might be due to the different ethnic groups recruited [14].

## Conclusions

In conclusion, our results show several microbial signatures in both, the vaginal and urinary microbiome and concentrations of specific HMOs being correlated with preterm labor, short cervix and PTB. In the vaginal microbiome, we identified *Gardnerella sp*. being associated with preterm labor, short cervix and PTB. *Ureaplasma sp*. correlated with short cervix and PTB and *L. jensenii*, *L. gasseri* and *Flavobacterium* associated with preterm labor, while *L. crispatus* was correlated with PTB. In the urinary microbiome, we detected changes in similar taxa, *Gardnerella* was the only taxa associated with preterm labor, short cervix and PTB, while other microbial signatures were associated only with PTB, for example *Ureaplasma urealyticum*, or only with short cervix, *Finegoldia* and *L. gasseri*, or with both preterm labor and short cervix, *L. jensenii* and *Staphylococcus*. We observed strong correlations between sialylated HMOs, especially 3’SL, and short cervix, preterm delivery and inflammation.

To sum up, our observations point at two different mechanisms leading to preterm labor and PTB. One seems to be driven by sterile inflammation, characterized by increased sialylated HMO concentrations in serum, and the second mechanism appears to be indirect and microbiome-mediated, which could, however, be driven by secretor-associated HMOs.

Our results identifying key HMOs and key microbial taxa associated with PTB will guide current efforts to better predict the risk for PTB in seemingly healthy pregnant women, and also provide appropriate preventive strategies.

## Materials and Methods

### Experimental Design and Clinical Metadata

60 pregnant women were recruited during a hospital visit due to potential preterm contractions. All study subjects had a viable pregnancy >23^+0^ weeks gestation, but not more than 34^+0^ weeks gestation, were healthy, were 18 years of age or older, willing to consent to all aspects of the protocol. Subjects were excluded if they had any recent genitourinary infections, multiple pregnancy, or more than 3 consecutive miscarriages, any known fetal anomalies associated with possible growth or genetic anomalies, subjects with pre-pregnancy diabetes type 1 or 2 or gestational diabetes mellitus, pre-pregnancy hypertension or if any antibiotic/prebiotic treatment was administered in the last 6 months. Clinical metadata from all subjects are listed in Suppl. Table 1. Gestational age is provided as weeks and days.

The grouping into cases (pregnant women who received tocolytic treatment) or controls (pregnant women who did not receive tocolytic treatment) was done based on medical examination. To diagnose preterm labor, subjects underwent cardiotocography to determine the uterine contractions and their frequency, and transvaginal ultrasonography for the measurement of the cervical length. Tocolysis was administered when ≥4 contractions / 30 minutes clearly felt by the patient were recorded by cardiotocography, and the patient had a short cervix (<25 mm). In Austrian hospitals, Atosiban, an oxytocin-receptor antagonist, is used as tocolytic treatment.

### Comments on the patient recruitment and classification into groups of high and low risk for preterm birth

Our study was originally designed to compare women with confirmed preterm labor (weeks 23-34) with gestational matched controls without contractions with regards to microbiome, HMOs and metabolites. We originally formed the groups of cases and controls (Fig. 7) on the basis of treatment with tocolytic agents following the diagnosis of preterm contractions. However, as the decision to administer tocolytic treatment remains to some extent subjective to the clinician in charge, using tocolysis treatment as proxy measure of preterm labor proved to be problematic. In our cohort, we had three women in the non tocolysis group with short cervix of whom two later delivered preterm. The tocolytic treatment was not administered to these two patients because they were close to 34 weeks of pregnancy (33^+6^ and 33^+5^) and it is not recommended to administer tocolytic treatment at 34 weeks of pregnancy [49]. Both patients received fetal lung maturation medication. Furthermore, one of these two patients had preterm premature rupture of the membranes (PPROM). The third patient who had a short cervix presented no signs of preterm contractions, consequently, did not receive tocolytic treatment and the outcome of the pregnancy was term delivery.

**Fig. 7.**
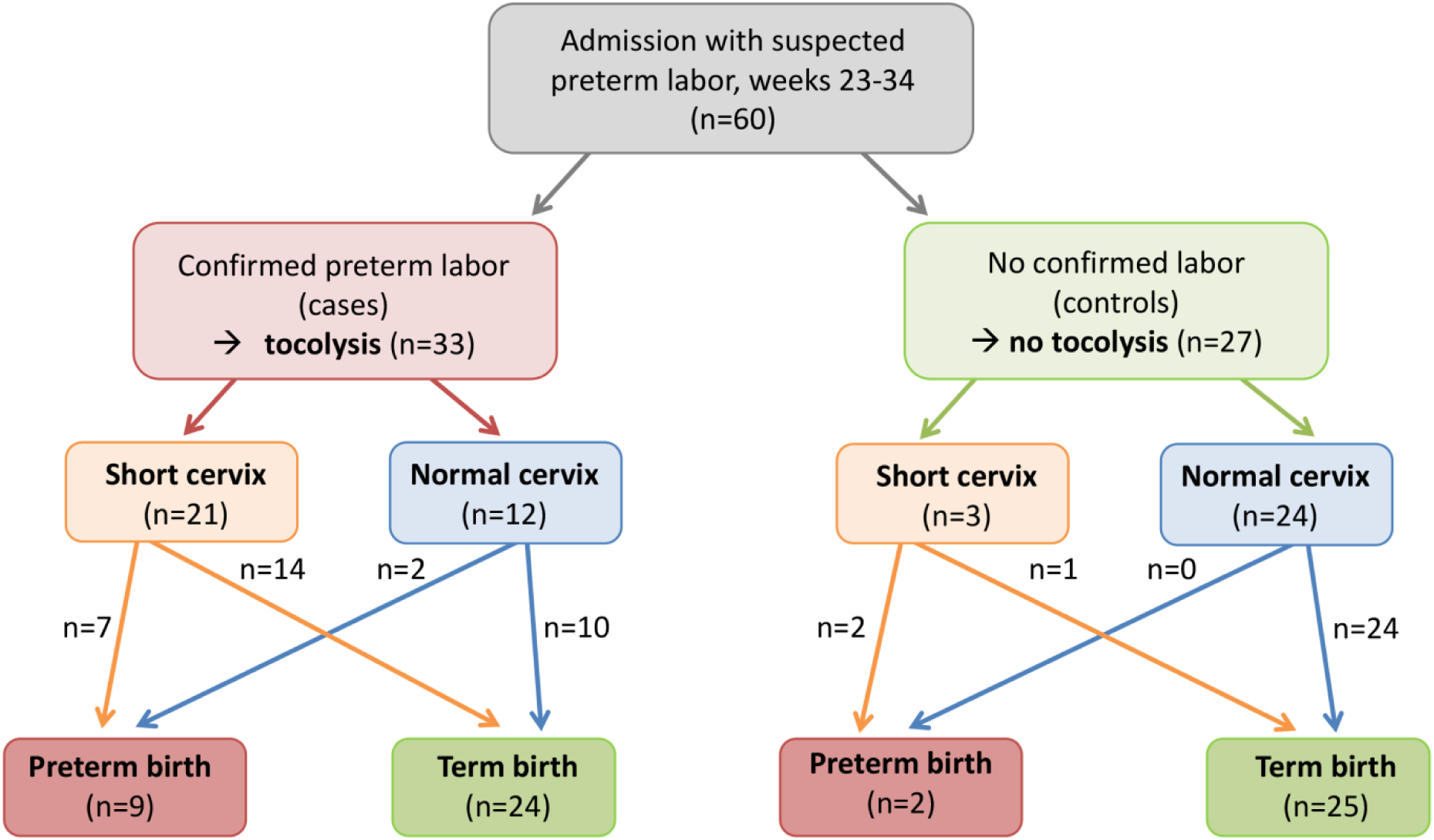
Flow chart study population

To add a more objective measure of risk for PTB, we also formed groups based on cervical length which has been previously associated with spontaneous PTB [49, 50].

In our cohort, 11 patients delivered preterm, 6 patients delivered before 34 weeks of pregnancy, and 5 patients delivered before 37 weeks of pregnancy. 9 of the women who delivered preterm had a short cervix (Fig. 7).

#### Sample Collection and Processing

Blood, urine and vaginal specimens were collected from each patient by trained personnel at the moment of recruitment and before medical treatment was administered. All urine specimens were collected using urinary catheters in sterile urine tubes. Vaginal swabs were collected using FLOQSwabs (Copan, Milan, Italy). All specimens were stored at 4°C and aliquots were prepared within 4h after collection and stored at −80°C until further processing.

Genomic DNA was extracted from the urine and vaginal specimens using the QIAamp DNA Mini Kit (QIAGEN) with some modification: before the extraction with the kit, the urine samples were centrifuged at 4400xg for 15 min, the supernatant was removed except for 500 µL that was used to resuspend the pellet. 500 µL of Lysis Buffer (sterile filtered, 20 mM Tris-HCl pH 8, 2 mM Na-EDTA, 1,2% Triton X-100) was added to the vaginal swabs. To both the vaginal and urine samples 50 µL of Lysozyme (10 mg/mL) and 6 µL of Mutanolysin (25 KU/mL) was added, followed by an incubation at 37°C for 1 h. The obtained mix was transferred to Lysing Matrix E tubes (MP Biomedicals) followed by a step of mechanical lysis at 5500 rpm for 30s two times using the MagNA Lyser Instrument (Roche, Mannheim, Germany). After the mechanical lysis the samples were centrifuged to separate the beads from the supernatant at 10000xg for 2 min. Afterwards the DNA was extracted according to the provided instructions. The DNA was eluted in 100 µL of Elution Buffer for the vaginal samples and in 60 µL for the urine samples. The genomic DNA concentration was measured using Qubit HS. Most urine samples had a DNA concentration under the detection limit.

#### PCR and quantitative PCR amplification

The genomic DNA was used to amplify the V4 region of the 16S rRNA gene using Illumina-tagged primers, 515FB and 806RB (Suppl. Table S2). The PCR reaction was performed in a final volume of 25 µL containing: TAKARA Ex Taq^®^ buffer with MgCl_2_ (10 X; Takara Bio Inc., Tokyo, Japan), primers 200 nM, dNTP mix 200 µM, TAKARA Ex Taq^®^ Polymerase 0.5 U, water (Lichrosolv^®^; Merck, Darmstadt, Germany), and DNA template. For vaginal samples 20-25 ng were used, while for urine 3-4 µL of genomic DNA was used. The PCR amplification conditions are listed in Suppl. Table S3. The bacterial 16S rRNA gene copies were determined using a SYBR based qPCR with the primer pair Bac331F-Bac797R (Suppl. Table S2). The reaction mix contained: 1x SsoAdvanced™ Universal SYBR^®^ Green Supermix (Bio-Rad, Hercules, USA), 300 nM of forward and reverse primer, gDNA template (20-25 ng for vaginal samples, 1 µL gDNA for urine samples), water (Lichrosolv^®^; Merck, Darmstadt, Germany). The qPCR was performed using the CFX96 Touch™ Real-Time PCR Detection System (Bio-Rad, Hercules, USA). The qPCR conditions used are given in Suppl. Table S3. All qPCR reactions were performed in triplicates. Crossing point (Cp) values were determined using the regression method within the Bio-Rad CFX Manager software version 3.1. Absolute copy numbers of bacterial 16S rRNA genes were calculated using the Cp values and the reaction efficiencies based on standard curves obtained from defined DNA samples from *Escherichia coli* [51]. The qPCR efficiency was between 90-105%, and the R^2^ values were always above 0.9. Detection limits were defined based on the average Cp values of non-template controls (triplicates) and the corresponding standard curves of the positive controls.

#### Amplicon sequencing, bioinformatics and statistical analysis

Library preparation and sequencing of the amplicons were carried out at the Core Facility Molecular Biology at the Center for Medical Research at the Medical University Graz, Austria. In brief, DNA concentrations were normalized using a SequalPrep™ normalization plate (Invitrogen), and each sample was indexed with a unique barcode sequence (8 cycles index PCR). After pooling of the indexed samples, a gel cut was carried out to purify the products of the index PCR. Sequencing was performed using the Illumina MiSeq device and MS-102-3003 MiSeq^®^ Reagent Kit v3-600cycles (2×251 cycles). The obtained MiSeq data is available in the European Nucleotide Archive under the study accession number: PRJEB31915.

The MiSeq data analysis was performed using QIIME2 [52] as described previously [53]. Shortly, after quality filtering, the DADA2 algorithm [54] was used to demultiplex, denoise truncated reads and generate ribosomal sequence variants (RSVs). The taxonomy was assigned using the SILVA v132 database [55]. The obtained feature table and taxonomy file were used for further analysis (Suppl. Table S4, S5).

The sequences classified as *Lactobacillus* genus were further used to classify to species level allowing the clustering of the vaginal microbiome into community state types based on the hierarchical clustering. The classification was performed through EzBioCloud [56]. To further test that the classification is correct a Maximum Likelihood tree was constructed in MEGA7 [57] using the sequences classified into the *Lactobacillus* genus. The alignment was performed in SINA [58], and the closes neighbors were picked only from the All-Species Living Tree project [59]. The tree can be found in the Suppl. Fig. S10.

The hierarchical clustering was performed in R on the relative abundance at species level based on Bray-Curtis dissimilarity and the method ward using the packages vegan [60] and gplots [61]. Alpha diversity indices were calculated using vegan package in R. Plots of the alpha diversity indices were generated in R using ggplot2 package [62]. Differences in the alpha diversity indices between the groups was tested in R using Wilcoxon Rank test. PCoA plots were constructed based on Bray-Curtis dissimilarities using phyloseq [63] and ggplot2 packages in R. The differences in the microbial composition between the analysed groups was tested in R using Adonis test. Barplots based on relative abundance at genera level were plotted for both the urinary and vaginal microbiome using ggplot2. LEfSe analysis plots and correlation heat maps were generated using Calypso [64]. Microbial functional profiles from 16S rRNA gene data were predicted by Tax4fun [65]. The results were visualized *via* Calypso. All statistical analyses were performed in SPSS (IBM Corp.) if not stated otherwise.

#### HMO isolation and analysis

Oligosaccharides from maternal serum were isolated as previously described [15]. In brief, 50 µL serum was diluted with H_2_O containing the internal standard Linear B6-Trisaccharide (Dextra Laboratories), and samples were subjected to 2 cycles of chloroform/methanol (2:1) extraction, followed by solid phase extraction (SPE) using C18 columns (Thermo Fisher Scientific) and graphitized carbon columns (Thermo Fisher Scientific). Urinary HMOs were isolated subjecting 10 µL urine samples with internal standard directly to C18 columns, then following the same protocol as for serum samples.

Isolated HMOs from serum and urine samples were fluorescently labeled with 2-aminobenzamide (2AB), as previously described [26]. The 2AB-glycans were separated by HPLC with fluorescence detection (360 nm/425 nm) on a TSKgel Amide-80 column (Tosoh Bioscience, Japan), using a linear gradient of a 50mmol/L-ammonium formate/acetonitrile solvent system. Commercially available standards for 2’-Fucosyllactose (2’FL), Lactodifucotetraose (LDFT), 3’-Sialyllactose (3’SL), 3’-Sialyllactosamine (3’SLN), and 6’-Sialyllactosamine (6’SLN) (Prozyme, Hayward CA) were used to annotate HPLC peaks. The amount of each individual HMO was calculated based on pre-determined response factors.

To normalize HMO concentration in urine samples, total osmolality indicative of solute concentration was measured in urine samples using an cryoscopic osmometer (Osmomat 030, Gonotec, Germany) according to the manufacturer’s protocol. HPLC Chromatogram area under the curve (AUC) for individual HMO peaks were divided by osmolality and multiplied by the mean of the osmolalities of all urine samples.

Secretor status was determined based on the relative abundance of 2’FL and LDFT.

HMO data are presented as median and IQR for skewed data. Changes in HMOs between groups were tested using Mann-Whitney-U-test, and graphs were plotted in GraphPad Prism (version 8.00; GraphPad Software, La Jolla California, USA).

#### Metabolomics

The phosphate buffer solution for NMR was prepared by dissolving 102 g of anhydrous KH_2_PO_4_ (VWR, Darmstadt, Germany), 0.5 g of TSP (3(trimethylsilyl) propionic acid-2,2,3,3-d_4_ sodium salt, Alfa Aesar, Karlsruhe, Germany), and 0.065 g NaN_3_ (VWR, Darmstadt, Germany), in 500 mL of D_2_O (Cambridge Isotopes laboratories (Tewksbury, MA)) and adjusted to pH 7.4 with 1M NaOH (VWR, Darmstadt, Germany) and HCl (VWR, Darmstadt, Germany). 300 µl of this NMR buffer in D_2_O were added to the samples containing 200 µl of urine and transferred to 5 mm NMR tubes. All NMR experiments were performed at 310K on an Avance Neo Bruker Ultrashield 600 MHz spectrometer equipped with a TXI probe head. The 1D CPMG (Carr-Purcell_Meiboom_Gill) pulse sequence (cpmgpr1d, 128 scans, 73728 points in F1, 11904.76 HZ spectral width, 128 transients, recycle delays 4s) with water suppression using pre-saturation, was used for ^1^H 1D NMR experiments. Bruker Topspin version 4.0.2 was used for NMR data acquisition. The spectra for all samples were automatically processed (exponential line broadening of 0.3 Hz), phased, and referenced using TSP at 0.0 ppm using Bruker Topspin 3.5 software (Bruker GmbH, Rheinstetten, Germany).

Integral regions for metabolites of interest corresponding to a certain number of protons and for external standard were defined. Concentrations of compounds were determined using Chenomx NMR Suite 8.3 using internal standard concentration. All quantified metabolites were normalized to original concentration of the urine.

For multivariate statistical analysis NMR data were imported to Matlab vR2014a (Mathworks, Natick, Massachusetts, United States), regions around the water and TSP signals excluded, and probabilistic quotient normalization was performed to correct for differences in sample metabolite dilution. MetaboAnalyst (4.0) was the performed to identify changes in metabolites in the urine samples.

### Strengths and limitations

A particular strength of our study is the innovative and comprehensive design of the study including different approaches (NGS for vaginal and urinary microbiome, urinary metabolome and measurement of HMOs in urine and blood) to determine and understand possible differences in pregnancy in women at high risk of preterm labor. While previous studies have focused on mostly vaginal microbiome or to a lesser extent, on the urinary microbiome, we here included both microbial communities. Introducing urinary HMOs in pregnancy will stimulate future studies investigating how HMOs can shape the urinary and vaginal microbiome. This study was designed as a cross-sectional pilot study, resulting in a limited sample size and consequently a small number of women who delivered preterm (11). This allowed forming groups of women with higher risk (based on confirmed preterm labor and cervical length) and investigate bivariate correlation. However, controlling for multiple confounders (secretor status, history of preterm births, miscarriages, BMI, smoking) using multivariate regression models is not applicable due to the small sample size. Moreover, future studies should also use longitudinal prospective pregnancy studies covering blood and microbiome sampling also in the first and second trimester. Furthermore, it would be interesting to monitor effects of the tocolysis treatment on HMOs and the microbiome.

## List of abbreviations

PTB: Preterm birth
CST: community state type
HMO: human milk oligosaccharide
2’FL: 2’-Fucosyllactose
LDFT: lactodifucotetraose
3’SL: 3’-sialyllactose
3’SLN: 3’-siallyllactosamine
6’SLN: 6’-siallyllactosamine
FUT2: fucosyltransferase-2
LEfSe: linear discriminant analysis effect size
PCoA: principal coordinate analysis

## Declarations

### Ethics approval

Research involving human material was performed in accordance with the Declaration of Helsinki and was approved by the local ethics committees (the Ethics Committee at the Medical University of Graz, Graz, Austria). The samples included in this study have been obtained covered by the ethics vote: 28-525 ex15/16.

### Consent of publication

Informed written consent was obtained from participants.

### Availability of data and materials

The obtained MiSeq data is available in the European Nucleotide Archive under the study accession number: PRJEB31915.

### Competing interests

All authors declare no competing interests.

### Funding

The research leading to these results has received funding from the People Programme (Marie Curie Actions) of the European Union’s Seventh Framework Programme FP7/2007-2013 under grant agreement number 627056 (to EJK). The study was financially supported by the City of Graz (to MRP) and by an anniversary fund of the Austrian National Bank (OeNB) under grant number 16927 (to EJK). MRP was trained in the Doctoral Program MolMed run at the Medical University Graz. The authors acknowledge the support of the ZMF Galaxy Team: Core Facility Computational Bioanalytics, Medical University of Graz, funded by the Austrian Federal Ministry of Education, Science and Research, Hochschulraum-Strukturmittel 2016 grant as part of BioTechMed Graz. This research was supported by the Austrian Science Fund (P28854, I3792, DK-MCD W1226 to TM), the President’s International Fellowship Initiative of CAS (No. 2015VBB045, to TM), the National Natural Science Foundation of China (No. 31450110423, to TM), the Austrian Research Promotion Agency (FFG: 864690, 870454), the Integrative Metabolism Research Center Graz, the Austrian infrastructure program 2016/2017, the Styrian government (Zukunftsfonds) and BioTechMed/Graz.

### Author contributions

MRP, EJK, VKK and CME designed the experiments and conceptualized the research. ECW and VKK recruited patients and collected clinical samples. MRP prepared, analyzed and interpreted the microbiome part of the study. CME performed the analysis, visualization and interpretation of the Tax4Fun results. EJK and EG performed HMO measurement and data analysis. GLR and TM performed metabolomics analysis and analyzed the metabolomics data. MRP, EJK and CME drafted the manuscript and interpreted the data. TM, ECW and VKK revised the manuscript for intellectual content. EJK and CME share primary responsibility for final content. All authors read and approved of the final manuscript.

## Acknowledgments

We acknowledge technical and advisory support provided by Bettina Amtmann, Petra Winkler, Birgit Freimüller and Karl Tamussino, Department of Obstetrics and Gynecology, Medical University of Graz.

